# PMTPred: Machine Learning Based Prediction of Protein Methyltransferases using the Composition of k-spaced Amino Acid Pairs

**DOI:** 10.1101/2023.09.20.558595

**Authors:** Arvind Kumar Yadav, Pradeep Kumar Gupta, Tiratha Raj Singh

**Author notes:** **Corresponding author:** Dr. Tiratha Raj Singh, Professor, Department of Biotechnology and Bioinformatics, JUIT, (H.P.), India, Mob No: 91-9816954695.

## Abstract

Protein methyltransferases (PMTs) are a group of enzymes that help to catalyze the transfer of a methyl group to its substrates. These enzymes play an important role in epigenetic regulation and are able to methylate various substrates with DNA, RNA, protein, and smallmolecule secondary metabolites. Dysregulation of methyltransferases is involved in different types of human cancers. However, in light of the well-recognized significance of PMTs, it becomes crucial to have reliable and fast methods for identifying these proteins. In the present work, we propose a machine-learning-based method for the identification of PMTs. Various sequence-based features were calculated and prediction models were develped using different machine-learning methods. A ten-fold cross-validation technique was used for model training. The SVM-based CKSAAP model gave the best prediction and achieved the highest accuracy of 87.94% with balance sensitivity (88.8%) and specificity (87.11%) with MCC of 0.759 and AUROC of 0.945. Also, SVM performed better than the compared deep learning algorithms for the prediction of PMTs. Finally, the best model was implemented in standalone software of PMTPred to facilitate the prediction of PMTs. The PMTPred achieved 86.50% prediction accuracy with 82.33% sensitivity, 90.67% specificity and ROC value 0.939 on the blind dataset. The standalone software of PMTPred is freely available for download at https://github.com/ArvindYadav7/PMTPred for research and academic use.

## 1. Introduction

Protein methyltransferases (PMTs) are enzymes that catalyze the transfer of a methyl group from the methyl donor S-adenosyl-L-methionine (SAM) to their substrates. These enzymes play a key role in epigenetic regulation and can methylate various substrates including DNA, RNA, protein, and small-molecule secondary metabolites. PMTs have mainly been classified into two major classes known as 1) protein arginine methyltransferases (PRMTs) that methylate arginine residues, and 2) protein lysine methyltransferases (PKMTs) that methylate lysine residues of their protein substrates. Dysregulation of PRMTs has been associated with multiple diseases, including cancers, pulmonary disorders, and cardiovascular disease [1]. PKMTs are well known to play significant roles in normal physiology and disease situations [2]. It is a critical and dynamic post-translational modification (PTM) that can control the protein function and stability [3]. PMTs play a crucial role in epigenetic regulation and transcriptional events through histone methylation and non-histone methylation at lysine or arginine residues. It has been reported that overexpression of PRMTs and PKMTs is associated with various types of human cancers [4, 5]. Thus, PMTs have emerged as promising and novel anti-cancer targets. Many drug development programs are focusing on the development of PMT-based small molecular inhibitors [6]. These molecule inhibitors have enormous importance in the interpretation of biological functions and disease mechanisms of targeted enzymes. Many inhibitors have been discovered based on PRMTs, and PKMTs [7–9]. Several PMT inhibitors have reached human clinical trials as investigative cancer therapeutics [9, 10]. PMTs also have potential applications in biotechnology, chemical biology, and synthetic biology to produce a variety of synthetic and natural compounds [11, 12].

In the light of the well-recognized role of PMTs in anti-cancer therapy, pharmacological and biotechnological applications, it becomes crucial to have superficial methods for the identification of PMTs. Many diverse PMT sequences are present, but most of these are assigned as hypothetical, putative, or probable functions based on sequence similarity. However, various existing experimental approaches for the identification of this protein are costly, time-consuming, labor-intensive, and require specialized equipment. Due to these obstacles, computational techniques for the prediction of various proteins emerged as a powerful alternative approach. Previously, some studies were conducted for the computational identification of methyltransferase reviewed by Petrossian and Clark [13]. Those studied used sequence-based, motif-based, and Hidden Markov Model (HMM) based search approaches. The methyltransferases have a topologically distinct family of proteins [13]. Thus, the similarity among primary sequences was found in only a small region of the protein. The search approaches based on sequence alignment are time-consuming and low-sensitive that can lead to problems in the prediction of low-similarity proteins.

In this situation, effective computational tools can be used in addition to experimental procedures to help get over their inherent constraints. Using machine learning techniques together with existing biological data can be an alternative method to predict the correct protein from sequence information. Due to their effectiveness, machine learningbased approaches for biological sequence function analysis and prediction have gained popularity in recent years. Various machine learning and deep learning algorithms have been widely used to solve classification problems [14–17]. A variety of machine learning-based methods have been developed for protein classification in recent years [18–24]. These methods utilized features derived from known sequence data to provide probable solutions for unknown sequence data based on the models trained in machine learning algorithms.

In the present study, we developed a sequence-based computational method using a machine learning technique that would be helpful for efficient and accurate prediction of PMTs. Different amino acid-based compositions and physicochemical-based features were used to train the classifiers to facilitate the method. Finally, the composition of k-spaced amino acid pairs (CKSAAP) feature gives the maximum prediction accuracy with the Support Vector Machine (SVM) classifier. Previously, CKSAAP has been used in various protein prediction problems associated with PTMs [25–29]. Therefore, we offered to utilize a 1-space CKSAAP feature to construct the model for the prediction of PMTs. The final model-based standalone tool named PMTPred is available for academic and research use. We hope that PMTPred could serve as a powerful tool for the identification of PMTs.

## 2. Methodology

### 2.1. Dataset preparation

The used dataset in the present study was extracted from the public database NCBI (https://www.ncbi.nlm.nih.gov/), which includes all protein methyltransferase. In this manner, 41146 methyltransferase sequences were retrieved from different organisms. Sequences having a length <50 amino acids were removed. Hypothetical, partial, putative, predicted, and sequences having non-amino acid characters were also removed and finally identified a total of 17280 methyltransferase protein sequences. Negative data is a prerequisite to develop a supervised machine-learning-based model. Therefore, to create a negative dataset, a total of 33061 sequences of non-methyltransferase (non-PMT) proteins were retrieved from the NCBI database. A similar protocol used for methyltransferase was followed and identified 3009 non-methyltransferase sequences. To obtain a non-redundant set of protein sequences CD-HIT program [30] was used at a 40% sequence identity threshold. Thus, CD-HIT results in 2862 methyltransferase and 3009 non-methyltransferase protein sequences. To construct a balanced dataset, we have randomly selected 2862 proteins out of 3009 non-methyltransferase proteins. Therefore final dataset has 2862 methyltransferase and 2862 non-methyltransferase proteins. Hereafter, we called instances of the final dataset a positive and negative dataset, respectively.

Further, to create the training set and test set, we followed the 80:20 ratios. The 80% of the total number of sequences were used as a training set and the remaining 20% as a test set from both positive and negative datasets respectively. Therefore, 2290 and 572 sequences were used in training and testing datasets respectively. An equal number of positive and negative sets in both training and testing data were used because machine learning can produce an unbiased result on a balanced dataset [31]. The flowchart of the used methodology is shown in Figure 1.

**Figure 1.**
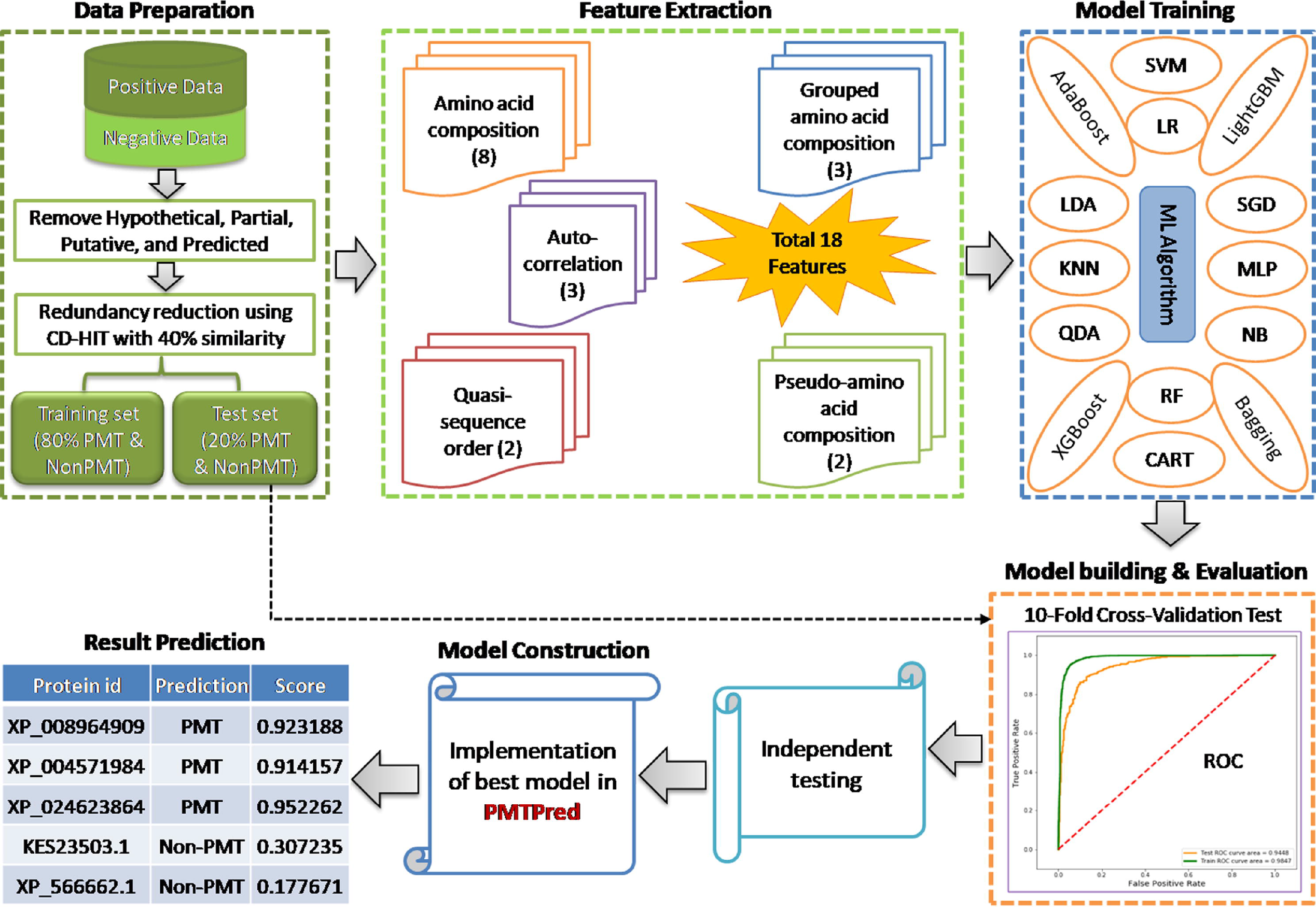
Flowchart of the used methodology in the present work. The main steps consisted of data preparation, feature extraction, model training, model building and evaluation, independent testing, and model construction.

### 2.2. Blind dataset

A blind dataset of PMTs was created from the UniProtKB database (https://www.uniprot.org/). We selected well-annotated PMT sequences from various organisms whose 3D structures were present. Sequence comparison analysis was performed through the CD-HIT-2D program [30] to ensure any similar sequence was not present in the main dataset. Then, randomly selected a set of 300 PMT sequences with a similar number of non-PMT sequences and created a blind data set to check the performance of PMTPred.

### 2.3. Feature extraction

To train the prediction model, various types of sequence-based feature descriptors were extracted. Different descriptor representation techniques have been used to convert protein sequences into numerical features. Feature extraction techniques are very important to build computational predictors [32–34]. In this study, we extracted five groups of descriptors such as amino acid composition (AAC), grouped amino acid composition (GAAC), autocorrelation, quasi-sequence-order (QSOrder), and pseudo-amino acid composition (PAAC). Each group may also include several feature extraction techniques. A total of 18 descriptors were calculated (Table 1) through a standalone Python-based toolkit of iFeature [35].

**Table 1.** Various feature descriptors and the number of descriptors in each group are calculated by iFeature.

| Descriptor group | Descriptor (abbreviation) | Number of attributes | Reference |
| --- | --- | --- | --- |
| **Amino acid composition** | Amino acid composition (AAC) | 20 | [36] |
| | Composition of $k$-spaced amino acid pairs (CKSAAP) | 2400 | [37, 38] |
|  | Dipeptides composition (DPC) | 400 | [36, 39] |
|  | Dipeptide deviation from expected mean (DDE) | 400 | [39] |
|  | Composition (CTDC) | 39 | [40–44] |
|  | Transition (CTDT) | 195 | [40–44] |
|  | Distribution (CTDD) | 39 | [40–44] |
|  | Conjoint triad (CTriad) | 343 | [45] |
| **Grouped amino acid composition** | Grouped amino acid composition (GAAC) | 5 | [33, 46] |
| | Composition of $k$-spaced amino acid group pairs (CKSAAGP) | 150 | [33, 46] |
|  | Grouped dipeptide composition (GDPC) | 25 | [33, 46] |
| **Autocorrelation** | Moran (Moran) | 240 | [47, 48] |
|  | Geary (Geary) | 240 | [49] |
|  | Normalized Moreau-Broto | 240 | [50] |
|  | (NMBroto) |  |  |
| **Quasi-sequence-order** | Sequence-order-coupling number (SOCNumber) | 60 | [51–53] |
|  | Quasi-sequence-order descriptors (QSOrder) | 100 | [51–53] |
| **Pseudo-amino acid composition** | Pseudo-amino acid composition (PAAC) | 50 | [54, 55] |
|  | Amphiphilic PAAC (APAAC) | 80 | [54, 55] |

In this study, the CKSAAP feature vector was used to represent the protein sequence in our proposed model. CKSAAP is an extensively used feature in bioinformatics problems [26, 56– 59]. It calculates the occurrence of amino acid pairs separated by any *k* number of amino acid residues. Here, *k* ranges were chosen from 0 to 5; meanwhile, CKSAAP gives the same result as DPC when *k* equals 0; therefore, *k* ranges from 1 to 5 are considered. For example, if *k*=0, the 0-spaced residue pair can be expressed as AA, AC, AD, …, YY (provide 400 amino acid residue pairs), and if *k*=1, then 1-spaced residue pair can be represented as AxA, AxC, AxD,…, YxY (provide 800 amino acid residue pairs). The CKSAAP feature is defined as:

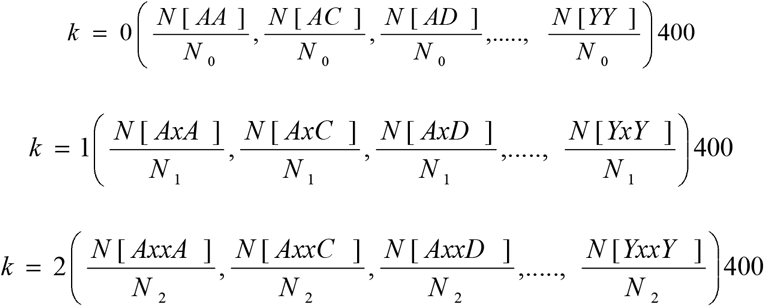

where ‘*x*’ stands for any of the 20 amino acids; N_k_ is calculated as *N*_*k*_= *L – (k + 1), k* = 1, 2, 3…, where *L* is the sequence length of a given protein. The final feature vector was calculated by concatenating the individual feature vectors. At *k* = 0, 1, 2, 3, 4, and 5, it generates a 2400-dimensional feature vector.

### 2.4 Machine-learning methods for model construction

Machine-learning approaches are the most effective methods for the development of prediction models, which are widely used in different fields. In this study, various machine-learning algorithms implemented with the Python library scikit-learn (Pedregosa et al., 2011) were trained to find out the best prediction model. A total of 14 machine-learning methods, namely Support Vector Machine (SVM) [61], Logistic Regression (LR) [62], Latent Dirichlet allocation (LDA) [63], k-Nearest Neighbors (KNN) [64], Classification and Regression Tree (CART) [65], NaiveBayes (NB) (Rennie et al., 2003), Random Forest (RF) [67], Multilayer Perceptron (MLP) (Rennie et al., 2003), Adaptive Boosting (AdaBoost) (Rojas, R., 2009), extreme gradient boosting (XGBoost) [69], light gradient boosting machine (LightGBM) [70], Stochastic Gradient Descent (SGD) [60], bootstrap aggregating (Bagging) [71], and Quadratic Discriminant Analysis (QDA) [63] were used. The grid search strategy was applied to find the best possible parameter combination using 10-fold cross-validation for each classification algorithm to obtain the best-performing model.

### 2.5 Model performance evaluation

The k-fold cross-validation technique was employed to measure the performance of classifiers. In this technique, each instance of the dataset is used to be tested once for prediction. Therefore, it is a purely unbiased method for the testing of model efficiency. In our study, we used a 10-fold cross-validation technique to measure the performance of classifiers. In this technique, the dataset was randomly divided into 10 equal subsets and one subset at a time was considered a test set, and the rest of the nine subsets were combined and used as a training set. This whole process was repeated 10 10-times so that each subset could be used as a test set at least one time. Finally, the scores of evaluation metrics of these 10 groups are averaged that evaluate the performance of the trained model. Then, an independent test set (data that is not present in the training set) was used to validate the performance of the models.

Further, to evaluate the performance of generated models, we have used various metrics like sensitivity (Sen), specificity (Spe), accuracy (Acc), precision (Pre), and Matthews correlation coefficient (MCC). All these parameters were computed by the utilization of the following formulas.

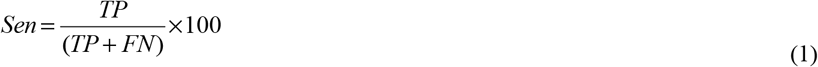

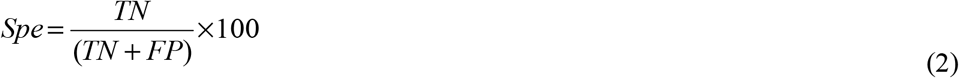

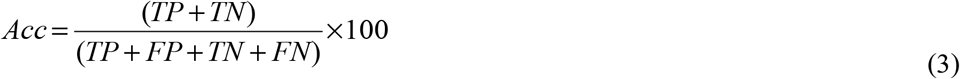

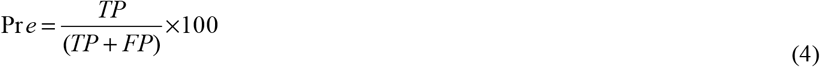

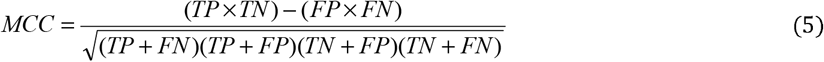

where, TP, TN, FP, and FN are the True Positives (correctly classified PMT), True Negatives (correctly classified non-PMT), False Positives (incorrect classification of non-PMT as PMT), and False Negatives (incorrect classification of PMT as non-PMT), respectively. Besides, we also calculated the area under the receiver operating characteristics (ROC) curve (AUROC) and the area under the precision-recall curve (AUPRC). The graphs for AUROC and AUPRC were plotted to estimate the visual assessment of the model. The ROC curve represents the true-positive rate vs. the false-positive rate, and the PR curve represents recall vs. precision. AUROC and AUPRC also presented the performance measure, and AUROC and AUPRC are close to 1, which signifies the best prediction of the model.

## 3. Result

## 3.1 Performance of different machine-learning algorithms

In this study, we have considered 18 feature calculation methods and calculated five different groups of descriptors. Each feature descriptor was trained with 14 machine-learning algorithms and measured the performance accuracy using a 10-fold cross-validation technique. Then the independent test dataset was applied to evaluate the performance of the models. Performance comparison analysis of all 14 methods using 18 features was done in terms of prediction accuracy (Supplementary Table S1). The best classification efficiency was achieved by 5-spaced CKSAAP feature with SVM classifier, followed by XGBoost, LightGBM, and MLP (Figure 2). SVM achieved prediction accuracy of 87.02 and 87.94% on the train and test set respectively. Therefore, based on performance, we find SVM as the best classifier for our dataset.

**Figure 2.**
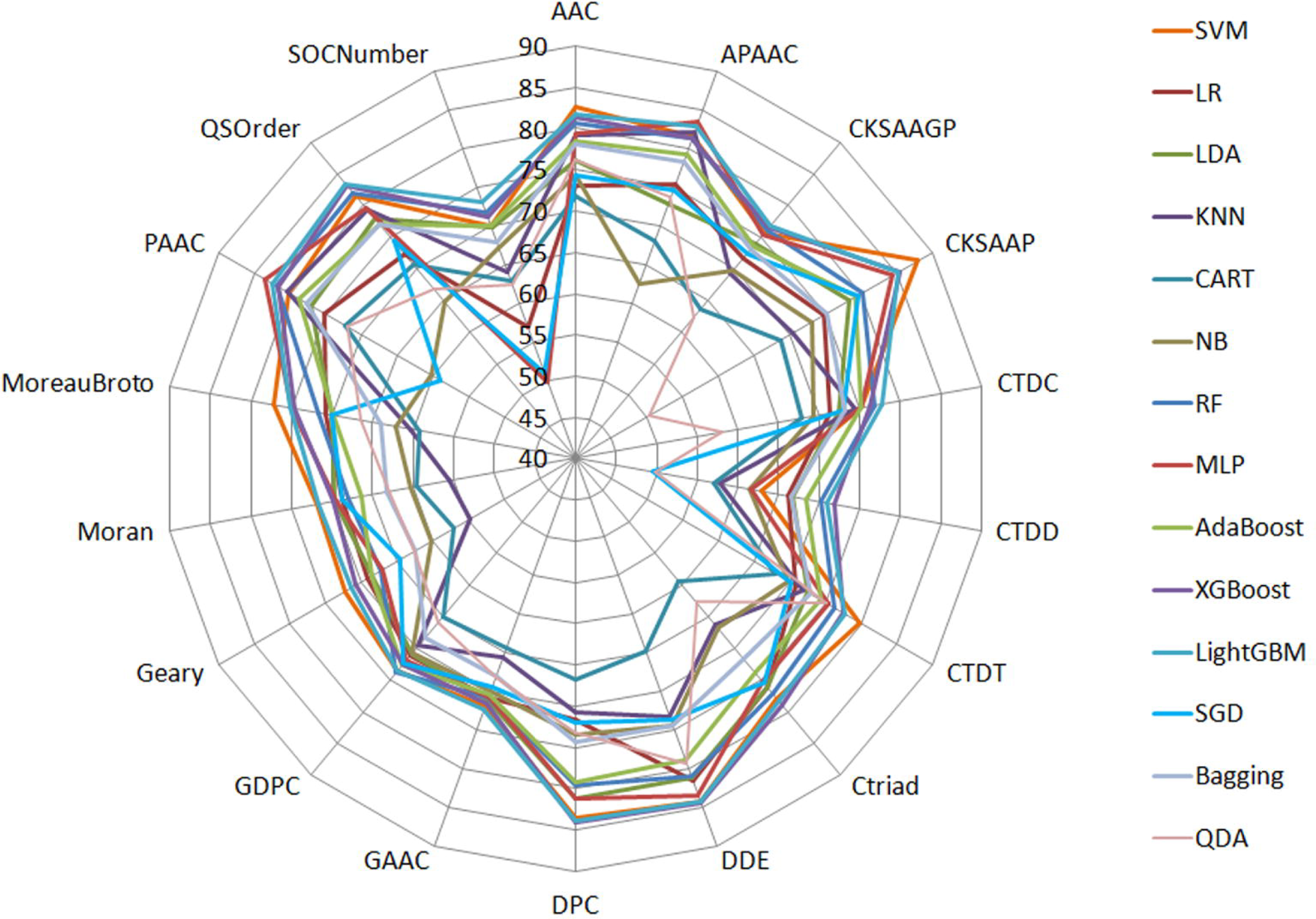
Radar chart representing the performance of different machine learning algorithms trained with various feature sets in terms of accuracy using the 10-fold cross-validation test. The different color lines represent various machine learning algorithms along with feature names on the outer line.

Further, to achieve the maximum accuracy, parameters tunning was performed for penalty parameter C (from 1.0 to 15.0 at step 1) and kernel parameter gamma (from 0.00001 to 5.0 at step 0.1) with radial basis function (RBF) kernel of SVM. SVM (at default parameters, C= 1 and gamma= scales) achieved maximum performance using CKSAAP with Sen of 89.38%, Spe of 88.48%, Acc of 87.94% with MCC of 0.759, and AUROC of 0.904. Although the AUROC of some features (DDE, DPC, and PAAC) was higher than CKSAAP, other metrics like Sen, Spe, Acc, and MCC were lower (Table 2). Our results revealed that SVM was a robust classification algorithm for the prediction of PMTs thus, SVM was adopted as a final classifier for this study.

**Table 2.** Top performance of each feature on test sets. For each row, we list the feature name, ML algorithm, and the number of features and performance evaluation of 10-fold crossvalidation.

| Feature | ML Algorithm | Performance evaluation |  |  |  |  |  |
| --- | --- | --- | --- | --- | --- | --- | --- |
|  |  | Acc | Sen | Spe | Pre | MCC | AUROC |
| AAC | SVM | 82.62 | 84.90 | 80.41 | 80.74 | 0.653 | 0.904 |
| APAAC | MLP | 83.39 | 81.17 | 77.49 | 77.72 | 0.664 | 0.877 |
| CKSAAGP | LightGBM | 76.85 | 78.68 | 75.08 | 75.34 | 0.538 | 0.841 |
| CKSAAP | SVM | 87.94 | 87.38 | 88.48 | 88.01 | 0.759 | 0.904 |
| CTDC | LightGBM | 77.70 | 78.89 | 76.50 | 77.41 | 0.554 | 0.847 |
| CTDD | XGBoost | 71.94 | 73.87 | 69.96 | 71.52 | 0.439 | 0.785 |
| CTDT | SVM | 79.80 | 85.64 | 73.85 | 76.98 | 0.599 | 0.859 |
| Ctriad | XGBoost | 79.02 | 77.16 | 80.91 | 80.50 | 0.581 | 0.864 |
| DDE | XGBoost | 84.45 | 84.36 | 84.53 | 84.07 | 0.689 | 0.912 |
| DPC | XGBoost | 84.10 | 85.61 | 82.64 | 82.67 | 0.683 | 0.923 |
| GAAC | SVM | 72.05 | 78.33 | 65.97 | 69.01 | 0.446 | 0.784 |
| GDPC | RF | 73.88 | 76.02 | 71.99 | 72.41 | 0.480 | 0.811 |
| Geary | SVM | 72.20 | 71.43 | 72.43 | 72.72 | 0.444 | 0.801 |
| Moran | SVM | 71.85 | 70.76 | 72.96 | 72.77 | 0.437 | 0.799 |
| MoreauBroto | SVM | 77.18 | 78.54 | 75.79 | 76.81 | 0.544 | 0.848 |
| PAAC | MLP | 83.49 | 85.96 | 83.67 | 83.59 | 0.696 | 0.912 |
| QSOrder | LightGBM | 83.40 | 84.36 | 82.47 | 82.32 | 0.668 | 0.908 |
| SOCNumber | LightGBM | 73.01 | 73.00 | 73.02 | 72.35 | 0.460 | 0.802 |

### 3.2. Performance of different CKSAAP feature subset

To obtain the optimum CKSAAP features with reduced vector size, we have evaluated the prediction performance on different CKSAAP feature subsets. Over-fitting and computational expense will be increased exponentially with the increase of k-value due to the large feature dimension. Thus, the dimension reduction of CKSAAP was carried out by changing the value of k-spaced. We have decreased the k-spaced value from 5 to 1 and reduced the dimensionality of the CKSAAP feature from 2400 to 800 attributes. Thus, four more feature sets with different attributes were further used to compare the prediction performance with 14 machine-learning algorithms using a 10-fold cross-validation technique. Out of 14 machine-learning methods, maximum performance accuracy was achieved by SVM for all CKSAAP feature subsets (Figure 3).

**Figure 3.**
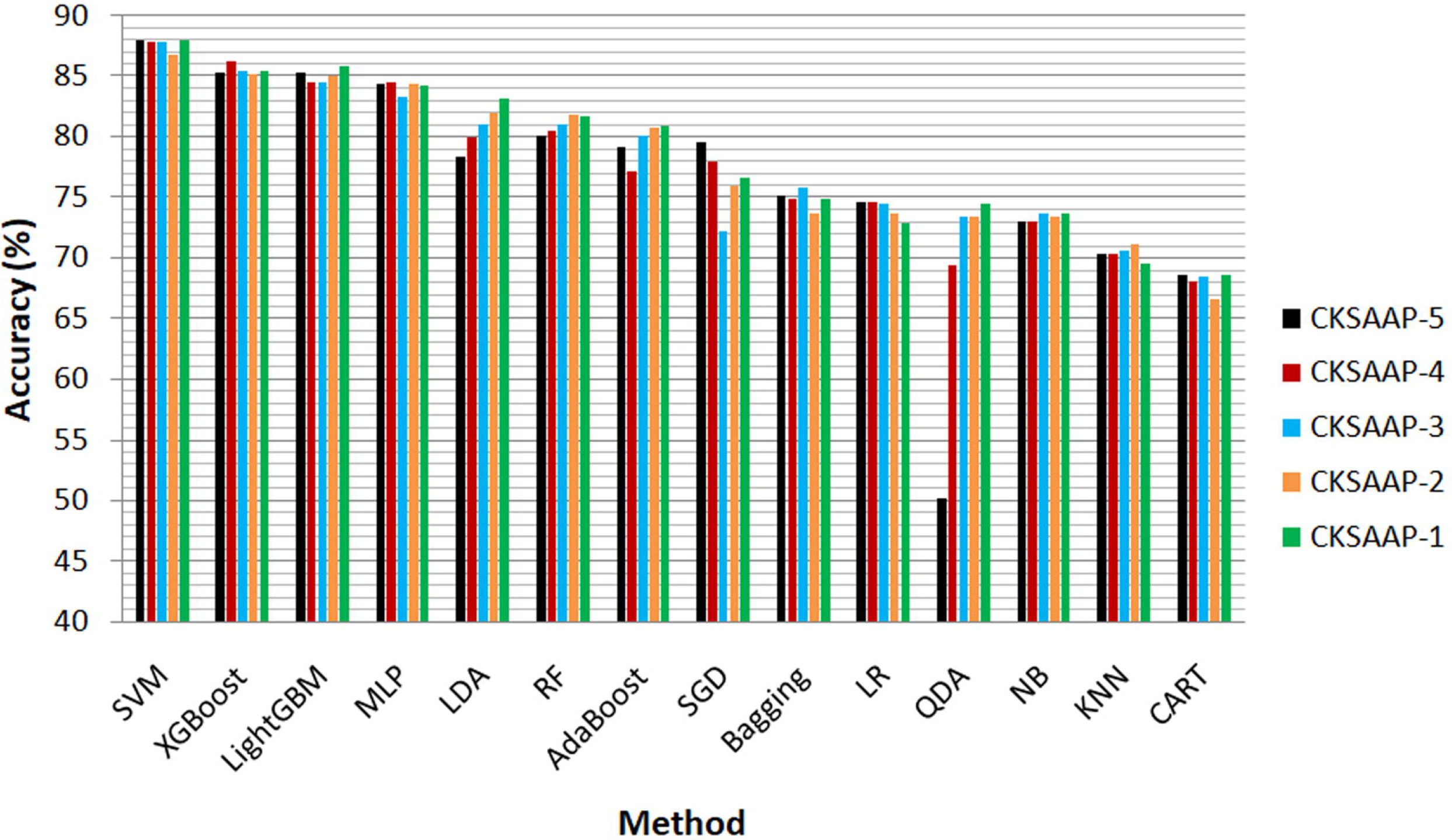
Performance comparisons of CKSAAP feature with a different set of descriptors using various machine-learning methods.

The accuracy performance of CKSAAP_k=1_ with vector size 800 was found similar to the performance of CKSAAP_k=5_ with vector size 2400. Both models of CKSAAP_k=5_ and CKSAAP_k=1_ achieved the maximum prediction accuracy of 87.94%. The maximum performance was achieved by SVM on c=1 and g= ‘scales’ in each feature set. CKSAAP_k=1_ has the highest Sen of 88.80% and MCC of 0.759 amongst all variants of CKSAAP with AUROC of 0.945. The accuracy of CKSAAP_k=4_ was almost similar to CKSAAP_k=1_ with an improved AUROC of 0.952 (Table 3).

**Table 3.** The feature selection process is driven by the performance of SVM on different kspaced values of CKSAAP.

| Feature | k-spaced value | Vector size | Acc | Sen | Spe | Pre | MCC | ROC |
| --- | --- | --- | --- | --- | --- | --- | --- | --- |
| CKSAAP<sub>k=5</sub> | 5 | 2400 | 87.94 | 87.38 | 88.48 | 88.01 | 0.759 | 0.904 |
| CKSAAP<sub>k=4</sub> | 4 | 2000 | 87.77 | 87.74 | 87.8 | 87.43 | 0.755 | 0.952 |
| CKSAAP<sub>k=3</sub> | 3 | 1600 | 87.86 | 88.27 | 87.45 | 87.19 | 0.757 | 0.949 |
| CKSAAP<sub>k=2</sub> | 2 | 1200 | 86.81 | 87.21 | 86.42 | 86.14 | 0.736 | 0.946 |
| CKSAAP<sub>k=1</sub> | 1 | 800 | 87.94 | 88.8 | 87.11 | 86.95 | 0.759 | 0.945 |

Figure 4A shows the performance of the CKSAAP_k=1_ model in terms of the ROC curve on training and testing datasets. The highest AUROC values achieved by the training and testing datasets model were 0.985 and 0.945, respectively. The model has an AUPRC value of 0.945 (Figure 4B). The ROC and PRC values towards one suggested that the CKSAAP_k=1_ model has better prediction ability.

**Figure 4.**
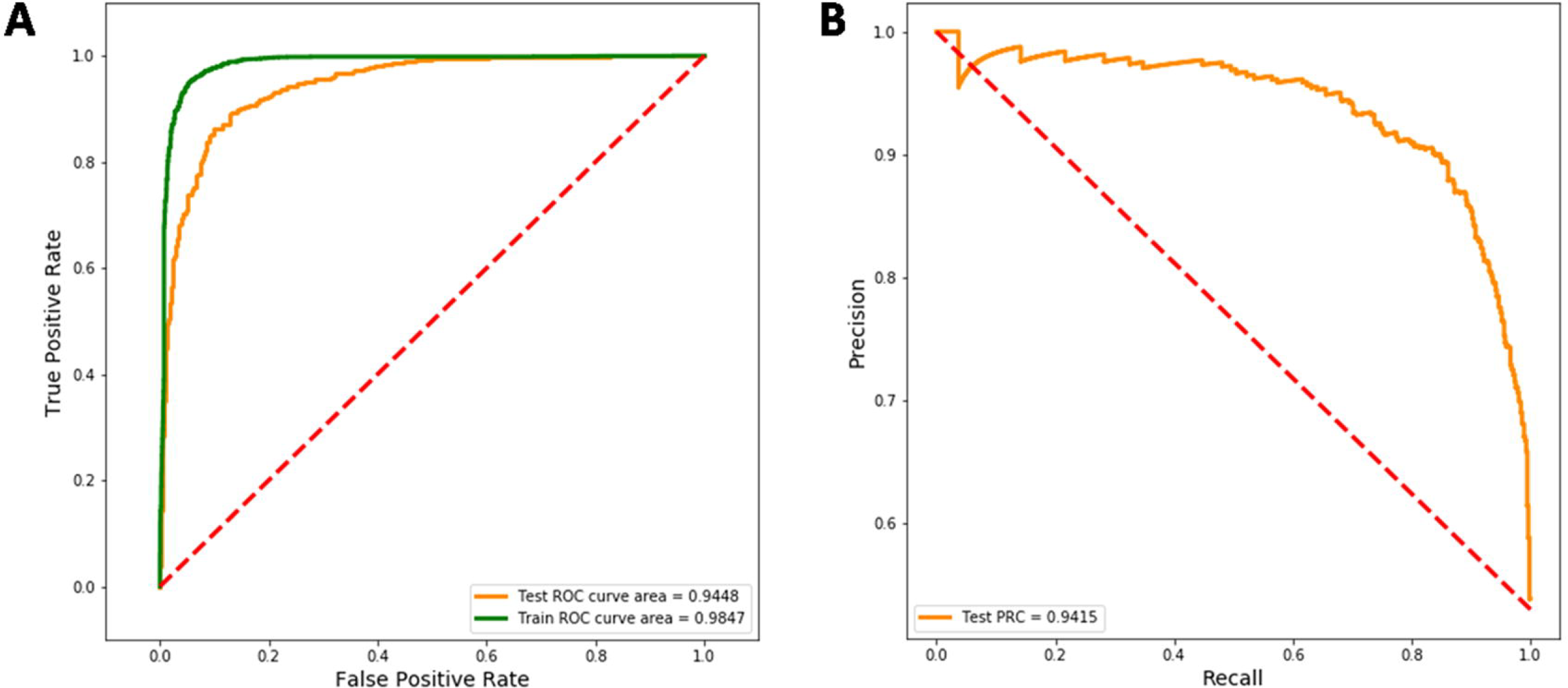
Graphs showing the performance of the CKSAAP_k=1_ model in terms of (A) Receiver operating characteristics (ROC) curve and (B) precision-recall curve (PRC).

#### Performance analysis of deep learning algorithms

Deep learning models such as convolutional neural networks (CNN) [72], attention-based convolutional neural network (ABCNN) [73] and Autoencoder (AE) [74] were also used to compare the performance with that of SVM. Deep learning was performed using the training data of the CKSAAP_k=1_ feature subset through 10-fold cross-validation. Among the deep learning models, AE performed well with the highest accuracy of 77.75% (Table 4). The CNN and ABCNN were not able to perform well. The performance metrics of deep learning models with all parameters are provided in Supplementary Table S2. The SVM was found to perform better than deep learning algorithms for PMT prediction as SVM outperformed the top-performing deep learning model AE by 10% in accuracy (Table 4). Therefore, the CKSAAP_k=1_ model based on best performance with a low-dimension feature vector was selected as a final model for the implementation purpose in our proposed PMTPred tool.

**Table 4:** Comparison metrics of SVM model with best performed deep learning model.

| Parameters | SVM | AE |
| --- | --- | --- |
| Accuracy | 87.94% | 77.75% |
| Sensitivity | 88.80% | 80.17% |
| Specificity | 87.11% | 75.33% |
| Precision | 86.95% | 76.30% |
| MCC | 0.759 | 0.563 |
| ROC | 0.945 | 0.859 |

### 3.3. Validation of PMTPred on a blind dataset

To evaluate the unbiased performance of the CKSAAP-based SVM classifier (PMTPred), a blind test evaluation has been carried out on the blind test dataset consisting of 300 PMT and 300 non-PMT protein sequences. The prediction accuracy of PMTPred was found to be 86.50% with 82.33% sensitivity, 90.67% specificity, 89.82% precision, 0.733 MCC, and 0.939 ROC. Hence, the blind-test performance established that CKSAAP-based SVM model has good prediction capacity.

## 4. Discussion

Growing evidence supports that, PMTs have been correlated with a variety of human malignancies. Nowadays, PMTs are a promising target class for therapeutic intervention in cancer [9, 75]. Thus, we have developed a machine-learning-based method for the prediction of PMTs called PMTPred. The relevance of PMTPred is to provide an easy-to-use prediction tool for methyltransferase-related proteins that have a prominent role in cancer progression and many other biotechnological products. The biological activity of any protein depends upon its basic sequence [76]; thus, we have used the sequence-based feature for model development. A total of 18 different numeric features were encoded using protein sequences. These numeric features were supplied to 14 machine-learning algorithms to train the models. This is because machine-learning methods are problem-specific [77]. Therefore, method selection requires exploring different methods on the same dataset for the selection of the best one. Model performance was evaluated to get the optimum model. All models were compared in terms of their prediction accuracy. It was observed that SVM performed well with the CKSAAP feature in comparison to other models (Supplementary Table S1). Due to good generalization ability, SVM has been effectively used in numerous classification problems [78–80]. Thus, the SVM-based CKSAAP model was selected as the final prediction model in this study. The CKSAAP feature has also been successfully used in various studies to address several prediction problems [18, 19, 24, 81, 82].

As shown in Table 2, the 5-spaced CKSAAP feature with 2400 attributes achieved the highest performance (accuracy of 87.94%) using SVM, followed by XGBoost, LightGBM, and MLP. The high-dimensional features eventually increase the computational complexity and may also contain irrelevant or redundant attributes that affect prediction accuracy [83, 84]. Therefore, to avoid the risk of over-fitting with a high-dimensional vector in SVM, we try to reduce the feature dimension of CKSAAP. For this purpose, more CKSAAP feature subsets were calculated with a decreased k-value from 5 to 1. These feature subsets were evaluated to find the best feature with a reduced dimension size (Figure 3). Maximum accuracy was achieved by SVM and the performance of CKSAAP at different k-spaced values did not change much. Performance assessment showed that CKSAAP_k=1_ has retained an accuracy of 87.94% only with 800 feature dimensions. CKSAAP_k=1_ has the highest Sen of 88.80% and MCC of 0.759 amongst all variants of CKSAAP with ROC of 0.945 (Table 3). The value of Sen is more considerable because it can improve the identification accuracy of the positive sample by reducing its scope. The highest ROC value was achieved by the CKSAAP_k=4_ model. However, in the case of the CKSAAP_k=1_ model, a little change in ROC may help us to reduce the computation and lower the risk of over-fitting due to the large feature dimension. Due to the small dimension size, this model also helps to speed up the prediction and improves the efficiency. The good sensitivity and precision capacity of the CKSAAP_k=1_ model are fine enough to classify the protein classes. The AUROC and AUPRC values also demonstrated that the SVM-based CKSAAP_k=1_ model had a better predictive performance. Additionally, the performance of SVM was also compared with three different deep learning algorithms, including AE, CNN, and ABCNN using the training dataset of the CKSAAP_k=1_ feature. SVM outperformed all three deep learning algorithms. The lower performance of deep learning algorithms may be due to the small data set. Therefore, we selected the SVM-based CKSAAP_k=1_ model based on the best performance with a low-dimension feature vector for the implementation in PMTPred. Further, we have used the independent blind dataset to validate the prediction performance of PMTPred. On the blind dataset, the prediction accuracy of PMTPred was 86.50% with 82.33% sensitivity and 90.67% specificity. Finally, to serve the scientific community, we have provided the standalone version of PMTPred.

### Standalone PMTPred

To serve the research and academic community, we have developed the standalone PMTPred tool in Python programming for both the Windows and Linux platforms. The standalone tool and other required files are available for download at https://github.com/ArvindYadav7/PMTPred. Users can provide input sequence files in FASTA format, and the PMTPred prediction result can be saved to a CSV file. The output of the CSV formatted file contains information about sequence ID, prediction, and probability score.

## Conclusion

In this work, a computational approach for constructing the machine learning-based model was proposed and successfully utilized for the prediction of PMTs. We have applied various machine learning and deep learning algorithms to different sequence-based features. Based on performance, it was concluded that SVM offers the best possible model for the prediction of PMTs. The 10-fold cross-validation method revealed that composition-based CKSAAP features gave top performance. A low-dimensional CKSAAP feature model with better performance has been obtained through the iteration of the k-spaced value. We implemented the best and low-dimensional CKSAAP_k=1_ model in the PMTPred tool for the efficient prediction of PMTs. We believe that PMTPred would be very useful for the prediction of PMTs to the scientific community and will have myriad applications.

## Declarations

### Ethics Approval and Consent to Participate

Not applicable.

