## Supplementary Table 2 for "PMTPred: Machine Learning Based Prediction of Protein Methyltransferases using the Composition of k-spaced Amino Acid Pairs"

Table S2: Performance analysis of deep learning algorithms using CKSAAP_k=1_ feature as training data.

|  | **Parameter** | **Sn** | **Sp** | **Pre** | **Acc** | **MCC** | **F1** | **AUROC** | **AUPRC** |
| --- | --- | --- | --- | --- | --- | --- | --- | --- | --- |
| **Autoencoder**  **(AE)** | Dropout rate= 0.5, Learning rate= 0.001, Epochs= 1000 | 80.175 | 75.327 | 76.301 | 77.752 | 0.5633 | 0.7777 | 0.8587 | 0.8216 |
| **Convolutional Neural**  **Network (CNN)** | Dropout rate= 0.5, Learning rate= 0.001, Epochs= 1000 | 0.0 | 100 | 0.0 | 50.0 | 0.0 | 0.0 | 0.5 | 0.75 |
| **Attention-based**  **Convolutional Neural**  **Network (ABCNN)** | Dropout rate= 0.5, Learning rate= 0.001, Epochs= 1000 | 0.0 | 100 | 0.0 | 50.0 | 0.0 | 0.0 | 0.5 | 0.75 |
